# IKK controls naive T cell survival by repressing RIPK1 dependent apoptosis and activation of NF-κB

**DOI:** 10.1101/2021.12.27.474254

**Authors:** Fiona Carty, Scott Layzell, Alessandro Barbarulo, Farjana Islam, Louise Webb, Benedict Seddon

**Affiliations:** Institute of Immunity and Transplantation, Division of Infection and Immunity, University College London, Royal Free Hospital, London, UK

## Abstract

The Inhibitor of Kappa B Kinase (IKK) complex is a critical regulator of NF-κB activation. In addition, IKK has recently been shown to repress RIPK1 dependent extrinsic cell death pathways by directly phosphorylating RIPK1. Our previous work shows that normal thymopoiesis relies on IKK exclusively for repression of TNF triggered cell death pathways, and that NF-κB activation by IKK is redundant for development. The role of these pathways in mature naive T cells has not previously been reported. Here, we show that, like thymocytes, naive peripheral T cells require continued IKK1/2 expression for survival. In contrast, however, cell loss is only partially prevented by blocking extrinsic cell death pathways by either deleting *Casp8* or inhibiting RIPK1 kinase activity. Inducible deletion of *Rela* in mature CD4^+^ T cells also results in a significant loss of naive CD4^+^ T cells and loss of IL7R expression, revealing an additional reliance upon NF-κB for long term survival of mature T cells. Together, these data show that IKK dependent survival of naive T cells depends upon both repression of extrinsic cell death pathways and activation of NF-κB survival programme.

**One sentence summary:** IKK regulates naive T cell survival

## Main text

### Introduction

The NF-κB family of transcription factors play critical roles in controlling development and function of many cell types (Bonizzi and Karin, 2004). Canonical NF-κB signalling is mediated by hetero or homodimers of p50, RELA and cREL family members that are sequestered in the cytoplasm by inhibitory proteins, the Inhibitors of kappa B (IκB) family and the related protein NFKB1. The key regulator of NF-κB dimer release is the inhibitor of kappa-B kinase (IKK) complex, a trimeric complex of two kinases, IKK1 (IKKα) and IKK2 (IKKβ), and a third regulatory component, NEMO (IKKγ). IKK phosphorylates IκB proteins, targeting them for degradation by the proteasome and releasing NF-κB dimers to enter the nucleus.

In mouse T cells, studies of tissue specific knockouts reveal complex overlapping and distinct roles for IKK and NF-κB in T cell biology. Ablation of the IKK complex, either by deletion of NEMO (Schmidt-Supprian et al., 2003), or combined loss of IKK1 and IKK2 subunits (Webb et al., 2016) results in a developmental arrest in single positive (SP) thymocytes at the immature HSA^hi^ stage. Similar developmental blocks are observed in mice lacking the upstream activator of IKK, TAK1 (Liu et al., 2006; Wan et al., 2006; Xing et al., 2016). In contrast, expression of NF-κB REL subunits is redundant for thymic development and generation of mature peripheral T cells (Oh et al., 2017; Webb et al., 2019). An explanation for this apparent contradiction comes from two key observations. First, the trigger for NF-κB activation via IKK in developing thymocytes is not TCR but TNF. TCR signalling activates IKK, and thus NF-κB, via a complex comprising CARD11, BCL10 and MALT1 proteins (the CBM complex) that is critical for T cell activation(Egawa et al., 2003; Hara et al., 2003; Ruefli-Brasse et al., 2003; Ruland et al., 2001; Ruland et al., 2003) but not required for thymocyte development (Schmidt-Supprian et al., 2004). In contrast, blockade of TNF signalling rescues development in IKK1/2 deficient thymocytes (Webb et al., 2016). Second, recent studies reveal that the IKK complex has two functions during TNF signalling in T cells -activating NF-κB and directly repressing cell death by inhibiting the serine threonine kinase, RIPK1. Ligation of TNFR1 causes recruitment of TRADD, TRAF2, and the serine/threonine kinase RIPK1. The ubiquitin ligases TRAF2, cellular inhibitor of apoptosis proteins (cIAPs) and the linear ubiquitin chain assembly complex (LUBAC), add ubiquitin chain modifications to themselves and RIPK1, creating a scaffold that allows recruitment and activation of the TAB/TAK and IKK complexes that in turn activate NF-κB. This is termed complex I (reviewed in (Annibaldi and Meier, 2018; Vandenabeele et al., 2010)). A failure to maintain the stability of this complex results in the formation of cell death inducing complexes. In the presence of IAP inhibitors, IKK inhibitors or TAK1 inhibitors (Dondelinger et al., 2013; Dondelinger et al., 2015; Wang et al., 2008), a complex composed of TRADD, FADD, CASPASE 8 and RIPK1 forms that induces apoptosis, a function dependent upon RIPK1 kinase activity (Annibaldi and Meier, 2018; Dondelinger et al., 2016; Ting and Bertrand, 2016). Phosphorylation of RIPK1 by IKK blocks RIPK1 kinase activity and therefore its capacity to induce apoptosis (Dondelinger et al., 2019; Dondelinger et al., 2015). In thymocytes, it is this function of IKK, and not NF-κB activation, that is critical for their survival and onward development and accounts for the phenotype observed in IKK deficiency(Webb et al., 2019).

Although the roles of IKK and NF-κB signalling during T cell development are now better understood, the function these signals play in maintenance of mature peripheral naive T cells remains unclear. Because of the block in thymic development in IKK-deficient mice, it is unknown whether IKK dependent repression of RIPK1 death signalling is active and necessary in mature T cells. Inducible ablation of IKK2 does not impact peripheral naive T cell survival (Silva et al., 2014), so redundancy with IKK1 may be sufficient to maintain IKK activity. Furthermore, although NF-κB is not required for development, its signalling is activated in mature SP thymocytes (Webb et al., 2016; Xing et al., 2016) and may therefore be important in mature peripheral T cells. One validated NF-κB gene target in thymocytes is *Il7r* (Miller et al., 2014; Silva et al., 2014). CD4^+^CD8^+^ DP thymocytes do not express IL-7R. Following successful positive selection, *Il7r* expression is induced in SPs by signals from TNFRSF members including TNFR1 and CD27, and dependent upon NF-κB. Thymic induction of *Il7r* is essential for long term survival of naive T cells(Silva et al., 2014). Whether basal NF-κB signalling is required to maintain expression of IL-7R is unclear. *In vivo*, inducible deletion of IKK2 does not affect T cell survival or IL7R expression(Silva et al., 2014). In contrast, re-expression of IL7R in cultured T cells does require NF-κB (Miller et al., 2014).

In the present study, we asked whether tonic IKK activity was required for survival of mature peripheral naive T cells. We tested this by the inducible ablation of genes for IKK1 and IKK2 in CD4^+^ T cells. Our study reveals a critical role for IKK expression and activity for long term survival of naive T cells. In contrast to the role of IKK in thymocytes, we found evidence that IKK dependent naive T cell survival was mediated by both repression of extrinsic cell death pathways and activation of NF-κB, revealing the shifting functions of IKK and NF-κB signalling during T cell ontogeny.

### Results

#### Expression of the IKK complex is required for naive T cell survival

In order to investigate whether the IKK complex is required for naive T cell survival, we generated mice in which the genes for IKK1 and/or IKK2 could be inducibly deleted in normal peripheral T cells of adult mice. *Chuk^fx/fx^ Ikbkb^fx/fx^ Rosa26^CreERT2^* (IKK1/2^iR26^) mice were generated that express a tamoxifen hormone-inducible CreER construct from the *Rosa26* locus. IKK signalling is required for normal homeostasis of numerous tissues (Li et al., 1999; Pasparakis et al., 2002; Schmidt-Supprian et al., 2000; Tanaka et al., 1999; Zaph et al., 2007). Therefore, since *Rosa26* is expressed ubiquitously throughout the body, we generated bone marrow chimeras by reconstituting irradiated *Rag1*^-/-^ hosts with T cell depleted bone marrow from IKK1/2^iR26^ donors, so that gene ablation was restricted to the haematopoietic system. Control chimeras were generated using *Rosa26^CreERT2^* donor bone marrow. Following reconstitution, tamoxifen was administered to all chimeras by intraperitoneal injection for five consecutive days. Spleens and lymph nodes were harvested at various time points after the first tamoxifen injection and analysed by flow cytometry. Following treatment, a substantial reduction in the number of both CD4^+^ and CD8^+^ naive T cell populations was observed (Fig. 1A) demonstrating a requirement for IKK expression in the haematopoietic compartment for persistence of naive CD4^+^ and CD8^+^ T cell populations.

**Figure 1.**
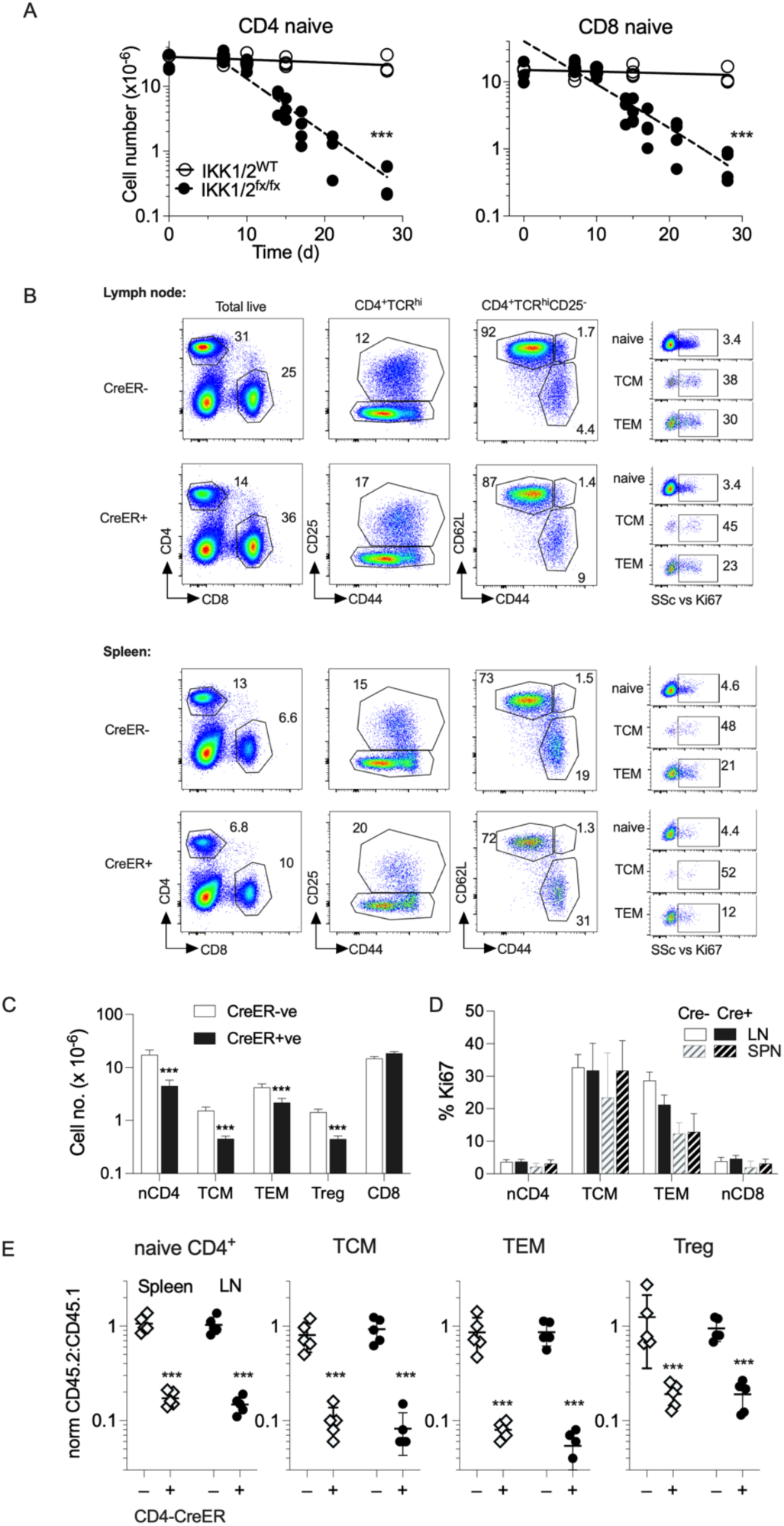
Inducible deletion of IKK1/2 genes cause a contraction in peripheral T cell numbers. (A) Bone marrow irradiation chimeras were generated by reconstituting sub lethally irradiated *Rag1^-/-^* mice with bone marrow from either IKK1/2^iR26^ or *Rosa26^CreERT^* donors (n=35/ group). 8 weeks later, hosts were treated with five injections of tamoxifen and mice culled at various times in the following 30 days. Scatter plots show the numbers of CD4^+^ and CD8^+^ naive T cells recovered from both LN and spleen of individual chimeras. Data are pooled from two independent experiments. Lines are best fit linear regressions for CreER -ve control groups and single phase decay kinetics for Cre+ mice. *** indicate significance of slope deviation from 0. (B-C) IKK1/2^iCD4^ and Cre-litter mates (n=10 and 8 respectively), were treated with tamoxifen for five days and analysed two weeks later. (B) Flow analysis of lymph node and spleen cells, shows the indicated phenotype of subpopulation gate indicated in the column for CreER+ and CreER-controls. Ki67 vs SSc was analysed for the indicated CD4^+^ subsets; naive (CD44^lo^ CD25^−^), T central memory (TCM, CD62L^hi^ CD44^hi^), T effector memory (TEM, CD62L^lo^, CD44^hi^). (C) Bar chart shows mean total cell recovery of the indicated subsets (nCD4, naive CD4; Treg, CD25^+^ regulatory T cells) from lymph node and spleen combined from the indicated hosts. (D) Bar chart shows mean % Ki67 expression by the indicated T cell populations from the indicated mice and organs. Data are pooled from three independent experiments. (E) Mixed bone marrow chimeras were generated by reconstituting lethally irradiated B6.SJL mice with 1:1 mix of bone marrow from (B6.SJL x C57Bl6/J)F1 donors and either CreER+ve or CreER-ve IKK1/2^iCD4^ donors. 8 weeks later, hosts were treated with five injections of tamoxifen and mice analysed a d21. Frequency of donor populations in lymph node and spleen was determined by flow analysis. Scatter plots show the fraction of donor cells derived from the indicated IKK1/2^iCD4^ partners for the indicated CD4 subsets, normalised to the donor fractions amongst CD8^+^ T cell compartment of the same mouse. Data are pooled from two independently generated batches of chimeras (n=5 total).

To specifically test whether IKK signalling was a cell intrinsic requirement for naive T cell survival, we bred mice in which IKK1 and IKK2 genes were inducibly deleted specifically in CD4 expressing T cells in *Chuk^fx/fx^Ikbkb^fx/fx^*CD4*^CreERT2^* (IKK1/2^iCD4^) mice respectively. Tamoxifen was administered for 5 days to IKK1/2^iCD4^ and Cre^−^ littermates and analysed two weeks after the last injection. Analysing phenotype of CD4 compartment in lymph node and spleen at this time revealed broadly normal representation of different CD4 subsets (Fig. 1B), but a clear reduction in the overall CD4 compartment size compared with CD8^+^ T cells, that were not targeted by Cre activity (Fig. 2B). Enumeration of different compartments confirmed that all CD4 naive, memory and regulatory compartments were specifically reduced following tamoxifen treatment, while CD8^+^ T cell numbers were similar (Fig. 1C). Although there was a loss of CD4^+^ T cells, we did not observed any evidence of lymphopenia or accompanying cell division, since Ki67 levels were similar in corresponding subsets from CreER^-^ and CreER^+^ hosts. Since we observed a loss across all compartments, we wished to confirm that the loss of naive CD4^+^ T cells was due to a cell intrinisic requirement for IKK expression, and not consequence of effects on other subsets. Therefore, we generated mixed bone marrow chimeras in which CD45.1 hosts were irradiated and reconstituted with equal mix of bone marrow from WT CD45.1/CD45.2 and either CD45.2 IKK1/2^iCD4^ mice or CreER -ve littermate donors. After eight weeks, all chimeras were treated with tamoxifen and two weeks later, the ratio of CD45.2:CD45.1 donor cells calculated for all CD4^+^ compartments. While normal ratios were maintained in chimeras reconstituted with CreER-bonemarrow, IKK deletion in progeny of IKK1/2^iCD4^ bonemarrow resulted in a profound loss in both naive and other CD4^+^ compartments.

**Figure 2.**
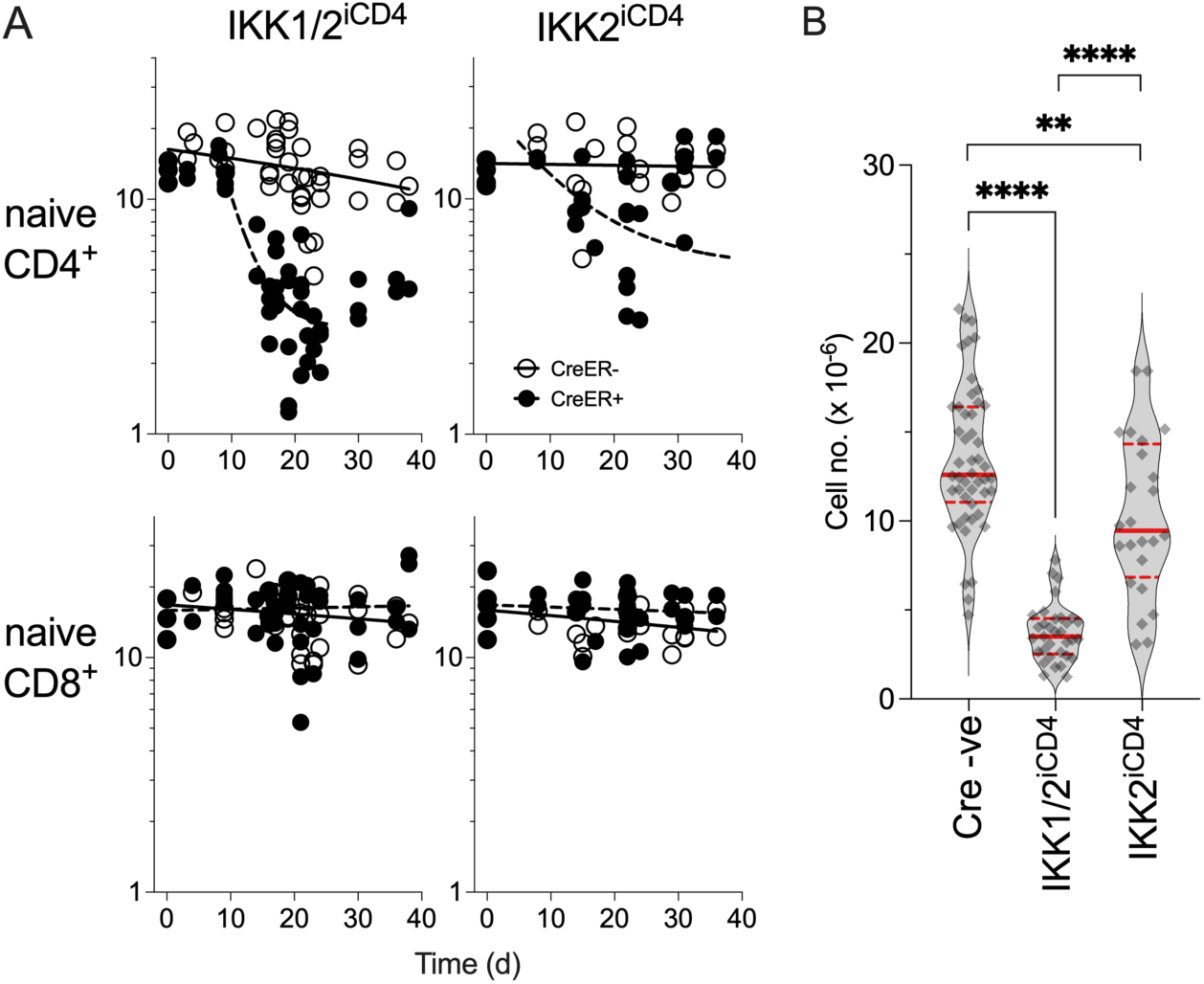
Kinetics of naive CD4^+^ T cell loss following IKK1/2 and IKK2 ablation. (A) Groups of IKK1/2^iCD4^ (n=46) and IKK2^iCD4^ (n=25) mice, together with Cre-litter mates (n=39 and 24 respectively), were treated with tamoxifen for five days. Mice from all groups were culled at various times following tamoxifen injections. Scatter plots show the numbers of CD4^+^ and CD8^+^ naive T cells recovered from both LN and spleen of individual mice. Line fits are best fit linear regressions for CreER -ve control groups and CD8^+^ naive T cells and single phase decay kinetics for CD4^+^ naive T cells in Cd4^CreER+^ mice. Data shown are the total pooled from 8 independent experiments. (B) Violin plot shows numbers of CD4^+^ naive T cells recovered from CreER -ve controls, IKK1/2^iCD4^ and IKK2^iCD4^ mice in (A), but binned from between days 10-30 post first tamoxifen injection.

To confirm gene deletion, we measured IKK1 and IKK2 protein in purified CD4^+^ naive T cells from tamoxifen treated mice immediately at d4, after the final injection or d21 (Fig. S1). In confirmation of our earlier work, we observed efficient deletion of IKK2(Silva et al., 2014). In contrast, ablation of IKK1 protein was reduced but still detectable. Inducible Cre mediated gene deletion is rarely completely efficient. So called ‘escapants’, that have failed to undergo gene deletion, are a common consequence in such systems, and are particularly evident when there is a selective pressure against successfully deleted cells, such as cell death. We therefore wanted to assess whether the observed cell loss was due to just IKK2 deletion or was a consequence of loosing both IKK1 and IKK2 gene expression. We therefore compared the impact of inducible deletion of IKK2, in *Ikbkb^fx/fx^*CD4*^CreERT2^* (IKK2^iCD4^) mice, with deletion of IKK1 and IKK2 in IKK1/2^iCD4^ mice over a range of time points following deletion. Inducible gene deletion did not influence the homing or distribution of naive T cells between lymph nodes and spleen (Fig. S2), and so we enumerated the total naive compartment sizes by combining total numbers in lymph nodes and spleen. Consistent with earlier reports(Silva et al., 2014), we observed only a modest and transient reduction in CD4^+^ naive T cell numbers following IKK2 deletion (Fig. 2A). In contrast, deletion of both IKK1 and IKK2 resulted in a more profound and sustained loss of naive CD4^+^ T cells numbers (Fig. 2A). As expected, the number of naive CD8^+^ T cells in the periphery remained stable over time in both strains (Fig. 2A). Statistical comparison of naive CD4^+^ compartment sizes, binning mice between d10 and d30, confirmed highly significant losses in IKK1/2^iCD4^ mice as compared with both CreER-controls and IKK2^iCD4^ mice (Fig. 2B).

#### IKK inhibition *in vitro* sensitises peripheral naive T cells to TNF induced cell death

In thymocytes, the IKK complex can both trigger NF-κB activation and protect cells from TNF induced cell death by direct phosphorylation and repression of RIPK1 kinase activity (Webb et al 2019). To test the acute function of IKK *in vitro*, we first asked whether IKK can also repress TNF induced cell death in mature naive T cells. We assessed the capacity of TNF to induce cell death when the IKK complex and RIPK1 activity was blocked using a combination of gene knockouts and kinase inhibitors. First, we analysed IKK1 deficient T cells from *huCD2*^iCre^ *Chuk^fx/fx^* (IKK1^ΔCD2^) mice cultured with IKK2 inhibitor (IKK2i), Bl605906 (Clark et al., 2011). T cells from Cre^−^ and Cre^+^ mice were cultured in increasing concentrations of recombinant TNF with or without optimal IKK2i, and the RIPK1 kinase inhibitor necrostatin-1 (Nec1), for 24 hours (Fig. 3A). Blocking mAbs against TNF and TNFR1 were also used in control wells to eliminate background cell death induced by TNF originating from cells within the culture. Cell death was induced at the highest concentrations of TNF in Cre^−^ control cultures treated with IKK2i (Fig. 3A). However, IKK1^ΔCD2^ T cells cultured with IKK2i were considerably more sensitive to TNF induced cell death, which was apparent even at the lowest concentrations of TNF (Fig. 3A). Cell death could be inhibited by the RIPK1 inhibitor Nec1, even at high TNF doses, suggesting that TNF induced naive T cell death following acute IKK blockade was completely dependent on RIPK1 kinase activity. We validated these findings in two additional ways. An alternative panIKK inhibitor, IKK16, rendered naive T cells susceptible to TNF induced cell death that could be blocked by Nec1 (Fig. 2B). Similarly, TNF induced cell death in the presence of IKK16 was prevented in T cells from mice expressing a kinase dead RIPK1^D138N^ mutant (Fig. 2C).

**Figure 3.**
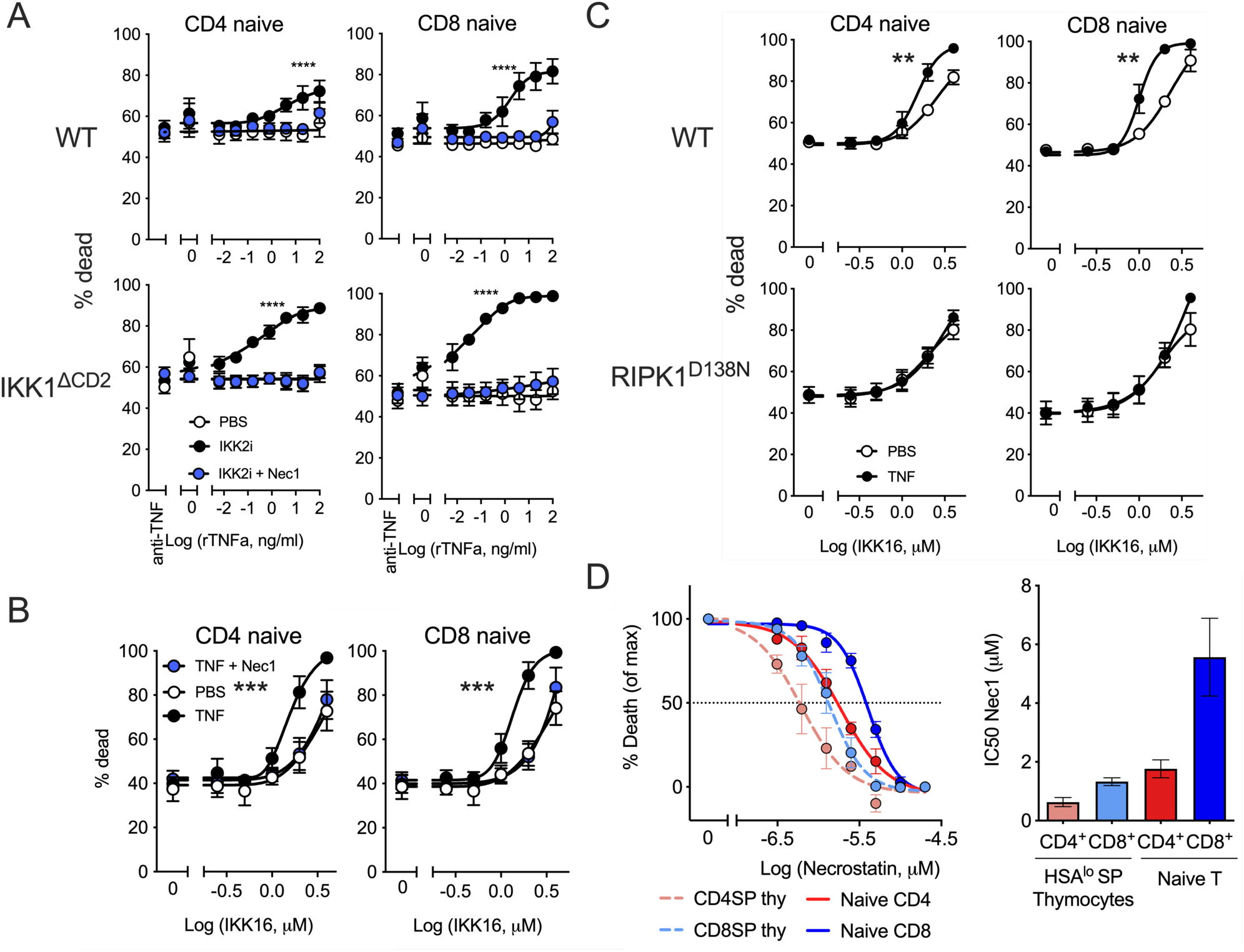
IKK blockade sensitises naive T cells to RIPK1 dependent TNF induced cell death in vitro. (A) LN T cells from WT and IKK1^ΔCD2^ mice (rows) were cultured with different doses of TNF, PBS, or TNF/TNFR1 blocking mAb for 24h, in the presence or absence of IKK2i and Nec1 inhibitors. Cultures were analysed by flow cytometry to determine viability (% dead) of naive CD4^+^ and CD8^+^ T cells (columns). (B) LN T cells from WT mice were cultured with the indicated concentrations of IKK16 inhibitor in the presence or absence of TNF and Nec1 inhibitor. Cultures were analysed to determine viability (% dead) of naive CD4^+^ and CD8^+^ T cells. (C) LN T cells from WT or RIPK1^D138N^ mice were cultured with the indicated concentrations of IKK16 inhibitor in the presence or absence of TNF. Cultures were analysed to determine viability (% dead) of naive CD4^+^ and CD8^+^ T cells. (D) Thymocytes and LN cells from IKK1^ΔCD2^ mice were cultured with IKK2i, TNF and different doses of Nec1. Cultures were analysed to determine viability (% dead) of naive CD4^+^ and CD8^+^ T cells in LN, and viability of HSA^lo^CD4^+^ SP and HSA^lo^CD8 SP thymocytes. Graphs show mean viabilities amongst the indicated populations from four (A), five (B) or three (C and D) independent experiments. Error bars indicate SEM (A-C) or 95% CI for IC50 calculations (D). *** indicated significance by 1 way ANOVA comparing PBS vs IKK2i (A) or PBS vs TNF (B, C) stimulations.

The capacity of IKK2i to sensitise WT naive T cells to some level of TNF induced cell death (Fig. 2A) contrasted with earlier findings in thymocytes, that are resistant to TNF induced cell death with IKK2i (Webb et al., 2016). Thymocytes become progressively more sensitive to IKK inhibition as they mature, and more so in CD8^+^ than CD4^+^ SPs, reflected by increasing IC50 of Nec1 required to block cell death (Webb et al., 2016). To test whether TNF induced death of naive T cells in the presence of IKK2i reflected a heightened sensitivity to IKK inhibition, we determined IC50s of Nec1 for naive CD4^+^ and CD8^+^ T cells as compared with mature HSA^lo^ SP thymocytes. Thymocytes and lymph nodes from IKK1^ΔCD2^ mice were cultured with the IKK2i, TNF, and increasing concentrations of Nec1 for 24 hours (Fig. 2D). Consistent with previous findings, the IC50 of Nec1 was higher in CD8^+^ SP than mature CD4^+^ SP thymocytes. The IC50 of Nec1 in cultures of peripheral T cells was higher still than in either thymocyte subset, with naive CD8^+^ T cells requiring the highest concentrations of Nec1. Therefore, developing T cells become progressively more sensitive to RIPK1 induced cell death as they mature in the thymus and following thymic egress.

#### Extrinsic cell death pathways dependent upon RIPK1 partially account for loss of naive CD4^+^ T cells following IKK1/2 ablation

The sensitivity of peripheral T cells to TNF induced cell death in vitro implies that one function of IKK in vivo may be to promote naive T cell survival by repressing extrinsic cell death pathways. Indeed, our earlier work shows that TNF is a critical trigger for cell death of thymocytes in the absence of IKK expression in vivo (Webb et al., 2016). However, the role of the extrinsic cell death pathway in the absence of IKK expression has not been explicitly tested either in thymocytes or in peripheral T cells. Apoptosis triggered by extrinsic death pathways is exclusively initiated by the aspartate-specific cysteine protease, Caspase 8 (Tummers and Green, 2017). Therefore, to directly assess the contribution of extrinsic death pathways to apoptosis of IKK deficient T cells, we analysed mice in which IKK1/2 and *Casp8* genes were simultaneously inactivated. To confirm the efficacy of our approach, we first analysed mice in which IKK and *Casp8* genes were deleted early in thymic development, in *Chuk^fx/fx^ Ikbkb^fx/fx^ Casp8^fx/fx^ Rosa26^REYFP^* huCD2^iCre^ (IKK1/2/*Casp8*^ΔCD2^) strain mice. As previously reported(Schmidt-Supprian et al., 2003), in the absence of IKK expression in *Chuk^fx/fx^ Ikbkb^fx/fx^ Rosa26^REYFP^* huCD2^iCre^ mice (IKK1/2^ΔCD2^), thymic development almost completely arrested amongst mature CD62L^hi^ HSA^lo^ single positive (SP) thymocytes (Fig. 4A-B). The few peripheral T cells present represent gene deletion escapants, evidenced by the lack of *Rosa26^REYFP^* Cre reporter expression (Srinivas et al., 2001) (Fig. 4C). Consistent with early findings that TNFR1 inactivation largely rescues thymic development (Webb et al., 2016), we found that *Tnf* ablation also mediates a comparable rescue of thymic development in IKK1/2^ΔCD2^ *Tnf*^-/-^ mice (Fig. 4A-B). However, *Tnf* ablation achieved only a very low level of peripheral T cell repopulation (Fig. 4C-D). In comparison, co-deletion of IKK1/2 and *Casp8* genes restored SP thymocyte numbers to WT levels (Fig. 4A-B), and also resulted in a substantial restoration of peripheral T cell numbers (Fig. 4C-D).

**Figure 4.**
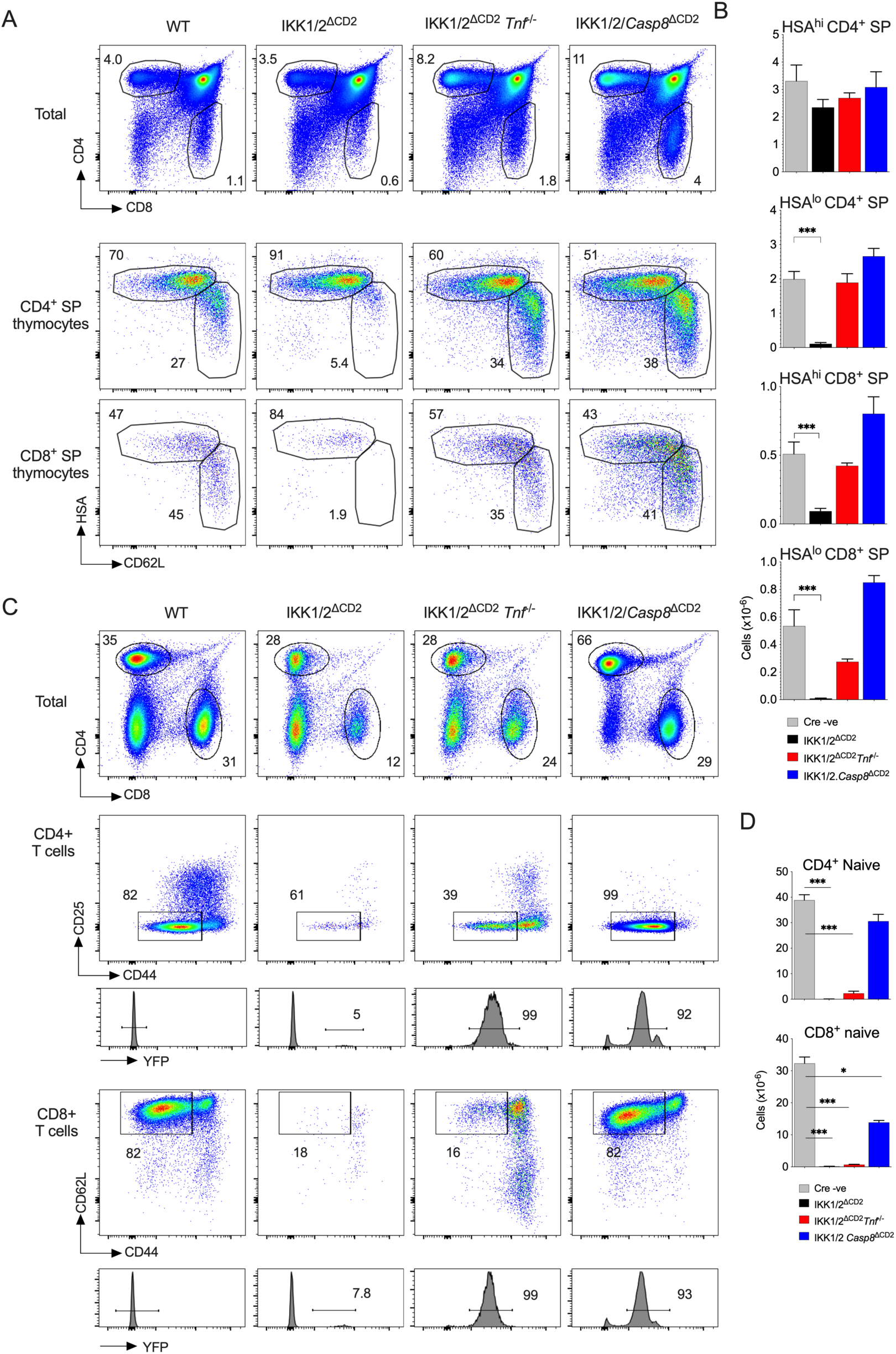
Casp8 deletion restores thymic development and peripheral repopulation following deletion of IKK1/2 genes in the T cell lineage. (A-B) Thymus from the indicated strains (n=4/strain pooled from two independent experiments) were enumerated and analysed by flow cytometry. Density plots are of CD4 vs CD8 for total live thymocytes (top row) or HSA vs CD62L for TCR^hi^ CD4^+^ SP and TCR^hi^ CD8^+^ SP thymocytes as indicated. Bar charts show total cell numbers of the indicated SP subsets for the indicated strains. (C-D) Lymph node and spleen were harvested from the same mice shown in A-B, enumerated and analysed by flow cytometry. Density plots are of CD4 vs CD8 for total live lymph node cells (top row), CD25 vs CD44 by TCR^hi^ CD4^+^ T cells (middle row) and CD62L vs CD44 by TCR^hi^ CD8^+^ T cells. Histograms are of Rosa26R^EYFP^ expression by naive T cells identified by the square gate in the corresponding density plot. Bar charts show total numbers of YFP^+^ naive T cells of the indicated lineage for the indicated strains recovered from both total lymph nodes and spleen. Error bars show SEM.

We next assessed the contribution of extrinsic cell death pathways of apoptosis to the loss of fully mature naive T cells following inducible IKK ablation. To do this, we measured the loss of CD4^+^ naive T cells in *Casp8^fx/fx^* IKK1/2^iCD4^ and *Casp8^fx^*^/WT^ IKK1/2^iCD4^ littermate controls, following induced gene deletion. Mice were treated with tamoxifen for five consecutive days, and then naive T cell numbers in lymphoid compartments quantified at day 22 after first injection. CD4^CreERT2^ induced IKK1/2 ablation in *Casp8^fx^*^/WT^ control mice resulted in a specific and substantial loss of CD4^+^ naive T cells (Fig 5A) comparable to that observed in *Casp8^WT^* IKK1/2^iCD4^ mice (Figs. 1 and 2). Complete ablation of *Casp8* alleles in IKK1/2^iCD4^ resulted in a partial rescue of CD4^+^ naive T cell numbers (Fig. 5B). Therefore, the requirement to repress extrinsic cell death pathways by IKK partially accounts for the loss naive CD4^+^ T cells following IKK1/2 ablation.

**Figure 5.**
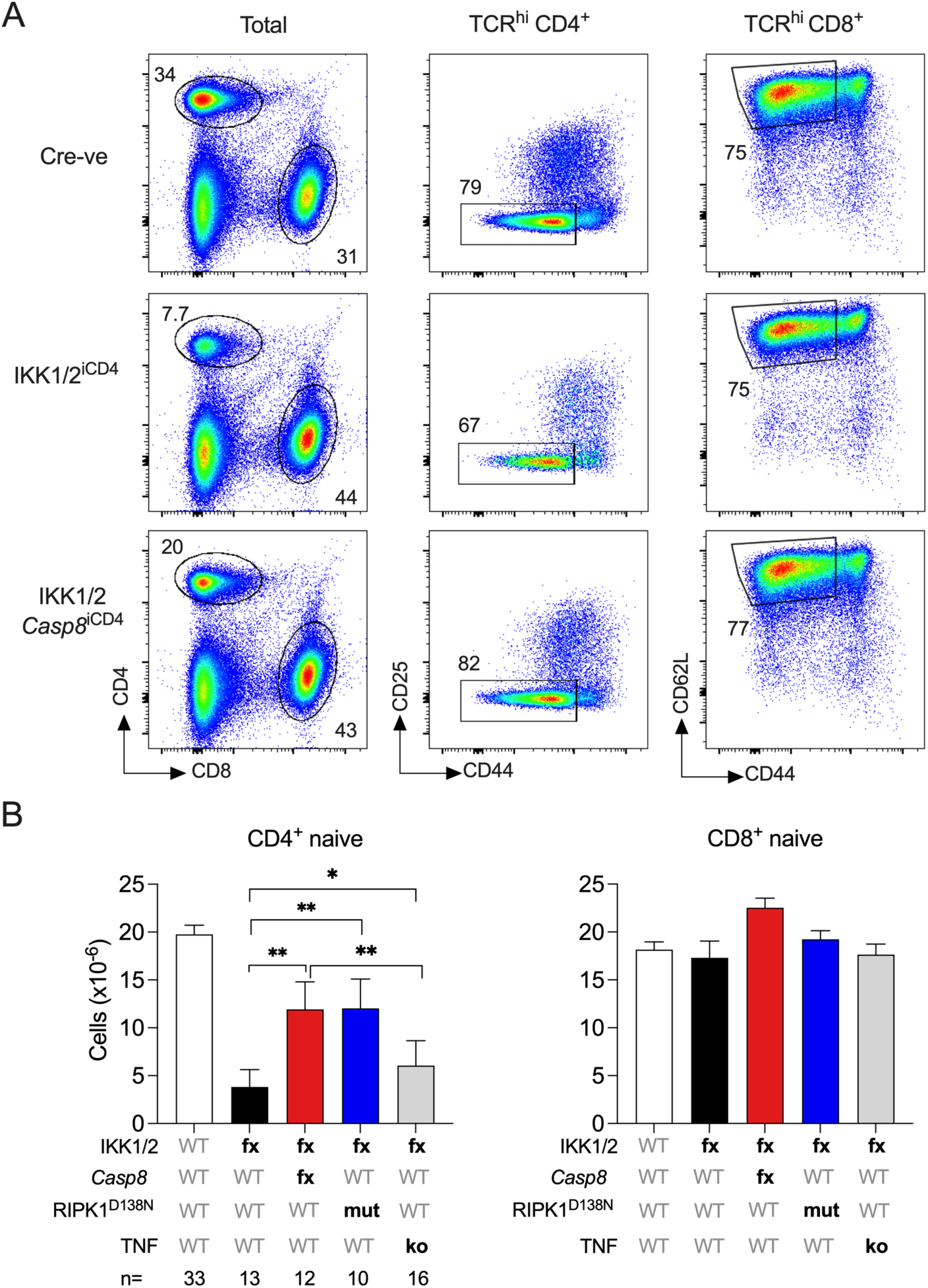
Inducible *Casp8* deletion or kinase dead RIPK1^D138N^ partially rescue naive T cells from IKK1/2 ablation in vivo. Groups of IKK1/2/*Casp8*^iCD4^, IKK1/2^iCD4^, IKK1/2^iCD4^ RIPK1^D138N^ and IKK1/2^iCD4^ TNF^KO^ mice, together with *Cd4^CreER^* -ve littermate controls were treated with tamoxifen for five days. At day 22, mice were culled and numbers of naive CD4^+^ and CD8^+^ T cells enumerated from LN and spleen. (A) Density plots show CD4 vs CD8 expression by total live lymph node cells, CD25 vs CD44 by TCR^hi^ CD4^+^ gated and CD62L vs CD44 by TCR^hi^ CD8^+^ T cells from lymph nodes. (B) Bar charts show total numbers of naive CD4^+^ and CD8^+^ T cells from the indicated mouse strains. Error bars indicate SEM. Data are pooled from five independent experiments.

The data from *in vitro* T cell cultures showed that IKK blocked RIPK1 dependent cell death in the face of extrinsic cell death stimuli such as TNF. In vivo, blocking extrinsic apoptotic cell death by ablating *Casp8* could rescue CD4^+^ naive T cells to about half normal levels, suggesting that one function of IKK in vivo is to repress CASPASE8 dependent cell death. We therefore wished to ask whether inhibition of CASPASE8 by IKK was mediated by the capacity of IKK to directly phosphorylate and repress RIPK1. To test this, we examined whether blocking RIPK1 kinase activity would achieve a similar rescue of naive CD4^+^ T cell numbers following IKK1/2 ablation as deletion of *Casp8*. We crossed IKK1/2^iCD4^ and kinase dead RIPK1^D138N^ (Newton et al., 2014) expressing strains in order to compare the effect of IKK1/2 deletion on naive CD4^+^ T cells with and without RIPK1 kinase activity. Tamoxifen was administered for five days to CreER^−^ and CreER^+^ IKK1/2^iCD4^ mice on both RIPK1^WT^ and RIPK1^D138N^ backgrounds, and lymph nodes and spleens were harvested on day 21 for analysis. By this time, deletion of IKK1/2 had caused a significant reduction in the number of naive CD4^+^ T cells of both RIPK1^WT^ and RIPK1^D138N^ mice, while CD8^+^ T cell numbers remained unaffected (Fig. 5B). While naive CD4^+^ T cells numbers dropped to less than 20% of control in IKK1/2^iCD4^ mice, numbers only dropped to ∼50% of control in IKK1/2^iCD4^ RIPK1^D138N^ mice. Therefore, blocking RIPK1 kinase activity achieved a similar rescue of naive CD4^+^ T cell numbers following IKK1/2 ablation as observed following corresponding ablation of *Casp8*, implying that CASPASE8 mediated cell death was indeed RIPK1 dependent.

We have previously shown that the block in thymic development observed in IKK1/2^ΔCD2^ mice can be largely overcome by inactivating TNFR1 signalling with null *Tnfrsf1a* alleles (Webb et al., 2016), a finding confirmed here using *Tnf*^-/-^ mice. In both cases, the rescue of development did not result in substantial repopulation of peripheral T cell compartments, which remained largely depleted. In contrast, *Casp8* deletion appeared to mediate a stronger rescue of peripheral T cell numbers in IKK1/2^ΔCD2^ than did blocking TNF signalling. These results suggests that TNF blockade alone is insufficient to rescue repopulation of naive compartments in IKK1/2^ΔCD2^ mice. However, to see if this was also true in established, mature naive compartments following induced deletion of IKK1/2, we generated *Tnf^-/-^* IKK1/2^iCD4^ mice. Consistent with observations in mice with constitutive T lineage targeted deletion of IKK1/2, *Tnf* deficiency only mediated a significant but small rescue naive CD4^+^ T cell numbers following inducible deletion of IKK1/2 (Fig. 5B). Taken together, these data suggest that RIPK1 is activated in naive T cells in the absence of IKK1/2 expression, but in contrast to thymocytes, by TNFR1 and additional receptors.

#### RELA expression is critical for survival of naive CD4^+^ T cells

Expression of canonical NF-κB subunits, RELA (p65), cREL and NFKB1, are not required for normal development of conventional αβ thymocytes (Webb et al., 2019). Evidence from here and previous work suggests that the exclusive survival function of IKK during T cell development is to repress RIPK1 dependent CASPASE8 triggered cell death. In contrast, data here showed that in peripheral naive CD4^+^ T cells, blocking extrinsic cell death pathways only partially protected the naive T cell compartment from the effects of inducible IKK1/2 deletion, suggesting that IKK serves additional functions for the survival of mature T cells. We therefore asked whether NF-κB activation downstream of IKK was important for naive T cell survival.

T cells express multiple REL family members and redundancy between these members results in complex interactions and phenotypes in the naive T cells, when one or more are constitutively deleted. Ablation of either cREL or RELA alone has no impact upon peripheral naive T cell numbers (Webb et al., 2019). Similarly, combined loss of cREL and NFKB1 has no effect on naive T cell numbers(Zheng et al., 2003). Together, these observations suggest that normal naive T cell numbers can be supported by either RELA alone, or by both cREL and p50, derived from NFKB1, in combination. Consistent with this view, combined deletion of RELA with either cREL or NFKB1 leads to a substantial reduction in T cell numbers (Webb et al., 2019). Therefore we tested whether mature naive CD4^+^ T cells require RELA for their survival by inducible ablation of *Rela* gene in *Rela^fx/fx^ Cd4^CreERT^* mice (*Rela^iCD4^*), in combination with either cREL or NFKB1 deficiency in *Rel*^-/-^ and *Nfkb1^-/^*^-^ strains or with *Rel^fx/fx^* alleles. Induced deletion of *Rela* following tamoxifen administration resulted in substantial loss of naive CD4^+^ T cell numbers by three weeks post-treatment, in both cREL deficient strains and *Rela^iCD4^ Nfkb1^-/-^* strains (Fig. 6A). Inducible deletion of *Rela* or *Rel* also impacted Treg survival (Fig. S3), as shown by others (Oh et al., 2017). Loss of naive CD4^+^ T cells occurred was independent of Treg or effects of gene deletion therein, since similar loss was observed in *Rel*^-/-^ *Rela^iCD4^* mice, that almost completely lack Treg (Fig. S3). Together, these data suggest that long term survival of naive CD4^+^ T cells does require tonic NF-κB signalling triggered by IKK activity.

**Figure 6.**
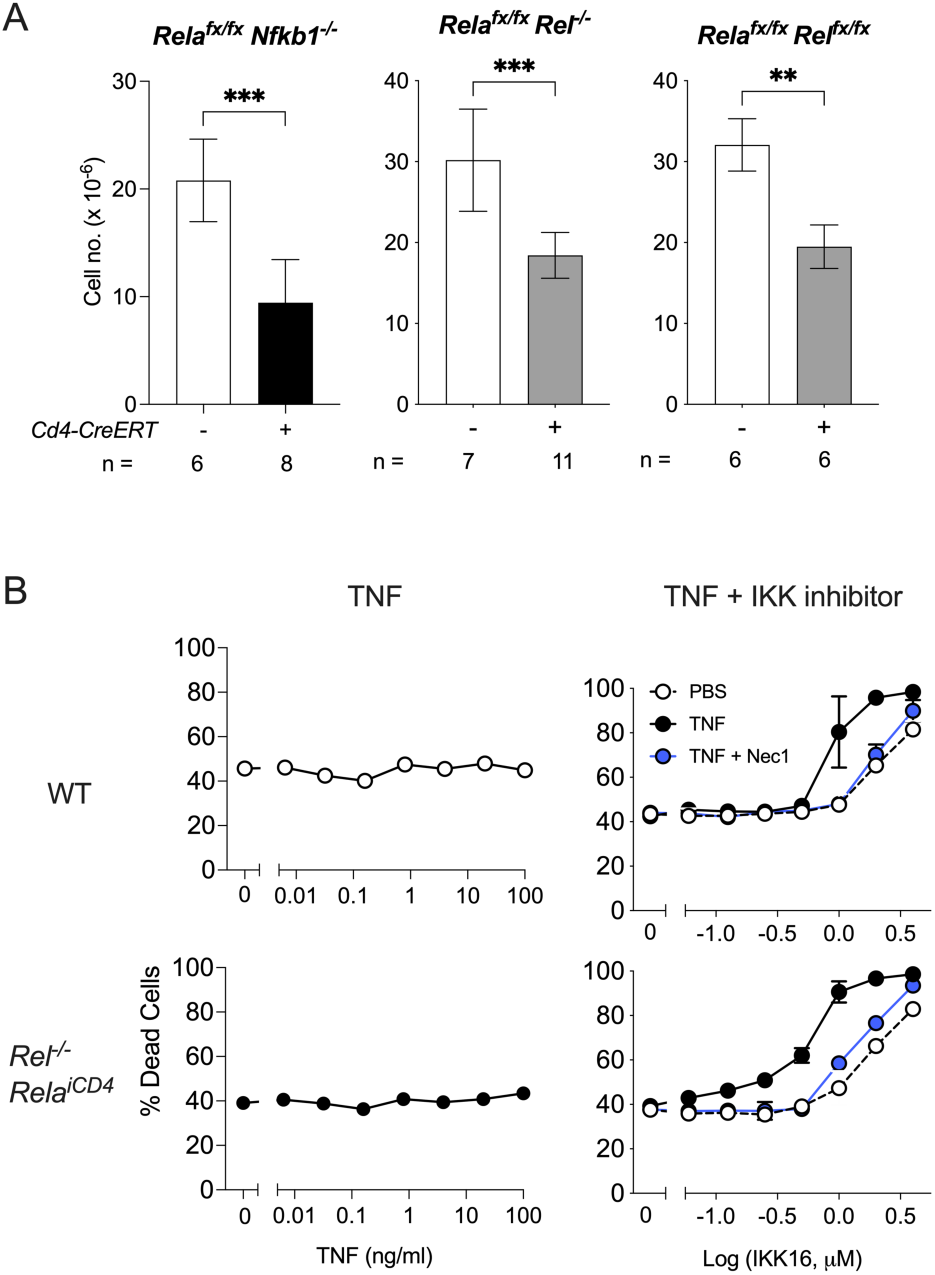
RELA expression is critical for survival of naive CD4^+^ T cells. Groups of *Rel^-/-^ Rela^iCD4^* (n=11), *Rela.Rel^iCD4^* (n=6) and *Nfkb1^-/-^ Rela^iCD4^* (n=8) mice, together with *Cd4^CreER^* -ve littermate controls (n=7, 5 and 6 respectively) were treated with tamoxifen for five days. At day 21, mice were culled and numbers of naive CD4^+^T cells enumerated from LN and spleen. (A) Bar charts show total numbers of naive CD4^+^ T cells of the indicated mouse strains (columns) from *Cd4^CreER^* +ve mice (filled bars) or *Cd4^CreER^ -*ve controls (empty bars). Error bars indicate SD. (B) LN cells from *Rel^-/-^ Rela^iCD4^* and WT mice in (A) recovered 21d following tamoxifen treatment were cultured for 24h with different concentrations of TNF, or with different concentrations of IKK16 inhibitor, with further addition of optimal concentrations of TNF or Nec1. Cultures were analysed by FACs to determine viability of CD4^+^CD44^lo^CD25^−^ naive T cells. Data are pool of five (A) or representative (B) of three independent experiments.

Our data suggest that IKK signalling promotes naive T cell survival both by repressing RIPK1-CASPASE8 triggered cell death and by activating an NF-κB dependent survival programme. Partial rescue of CD4^+^ naive T cell compartment following IKK1/2 deletion by *Casp8* ablation, implies that the dependence of naive T cells upon NF-κB signalling is independent of extrinsic cell death pathways. To confirm this, we asked whether naive CD4^+^ T cells remained resistant to cell death triggered by extrinsic cell death triggers, following tamoxifen induced ablation of *Rela*. We measured the sensitivity of RELA deficient CD4^+^ T cells to TNF induced cell death. Even when cultured with supra-physiological concentrations of TNF, cREL/RELA deficient T cells maintained viability in a comparable manner to WT controls (Fig. 6B). Survival of cREL/RELA deficient T cells was still dependent upon intact IKK1/2 signalling, since pharmacological inhibition with panIKK inhibitor, IKK16, sensitised both cREL/RELA deficient and WT CD4^+^ T cells to TNF dependent cell death, that could also be blocked by the RIPK1 inhibitor, Nec1. Therefore, the mechanism by which NF-κB promotes survival of naive CD4^+^ T cells appears independent of extrinsic cell death pathways.

#### NF-κB expression is required to maintain IL-7R expression in naive T cells *in vivo*

IL-7 signalling is a critical survival cue for T cells(Barata et al., 2019). Induction of *Il7r* in developing thymocytes is dependent upon NF-κB and is essential for the subsequent long term survival of naive T cells (Miller et al., 2014; Silva et al., 2014; Webb et al., 2019; Webb et al., 2016). Whether maintenance of *Il7r* expression in naive T cells also depends upon tonic NF-κB signalling is not known. To test this, we analysed IL-7R protein in both RelA^iCD4^ and IKK1/2^iCD4^ strains following inducible deletion of genes for RELA or IKK proteins. Protein levels of IL-7R were determined by flow cytometry and MFI of expression normalised to tamoxifen treated CreER^−^ controls analysed at the same time. We first analysed the kinetics of IL-7R expression by CD4+ naive T cells following deletion of IKK1/2 and IKK2 alone. Loss of IL-7R expression was observed soon after administration of tamoxifen in both IKK1/2^iCD4^ and IKK2^iCD4^ strains, reaching a maximal loss around d20 in both cases (Fig. 7A). Expression by CD8^+^ T cells remained normal. Analysing IL-7R expression at this three week time point post tamoxifen administration revealed comparable loss of IL-7R following RELA ablation, as well as in IKK1/2i^CD4^ strains with either *Casp8^fx/fx^* alleles or RIPK1^D138N^ mutations that rescued CD4^+^ T cells from extrinsic cell death (Fig. 7B). This confirmed that loss of IL-7R was independent of extrinsic death pathway activation. These data also show that constitutive expression of RELA is required to maintain IL-7R expression in naive T cells and that mature naive T cells are subject to tonic NF-κB signalling in vivo.

**Figure 7.**
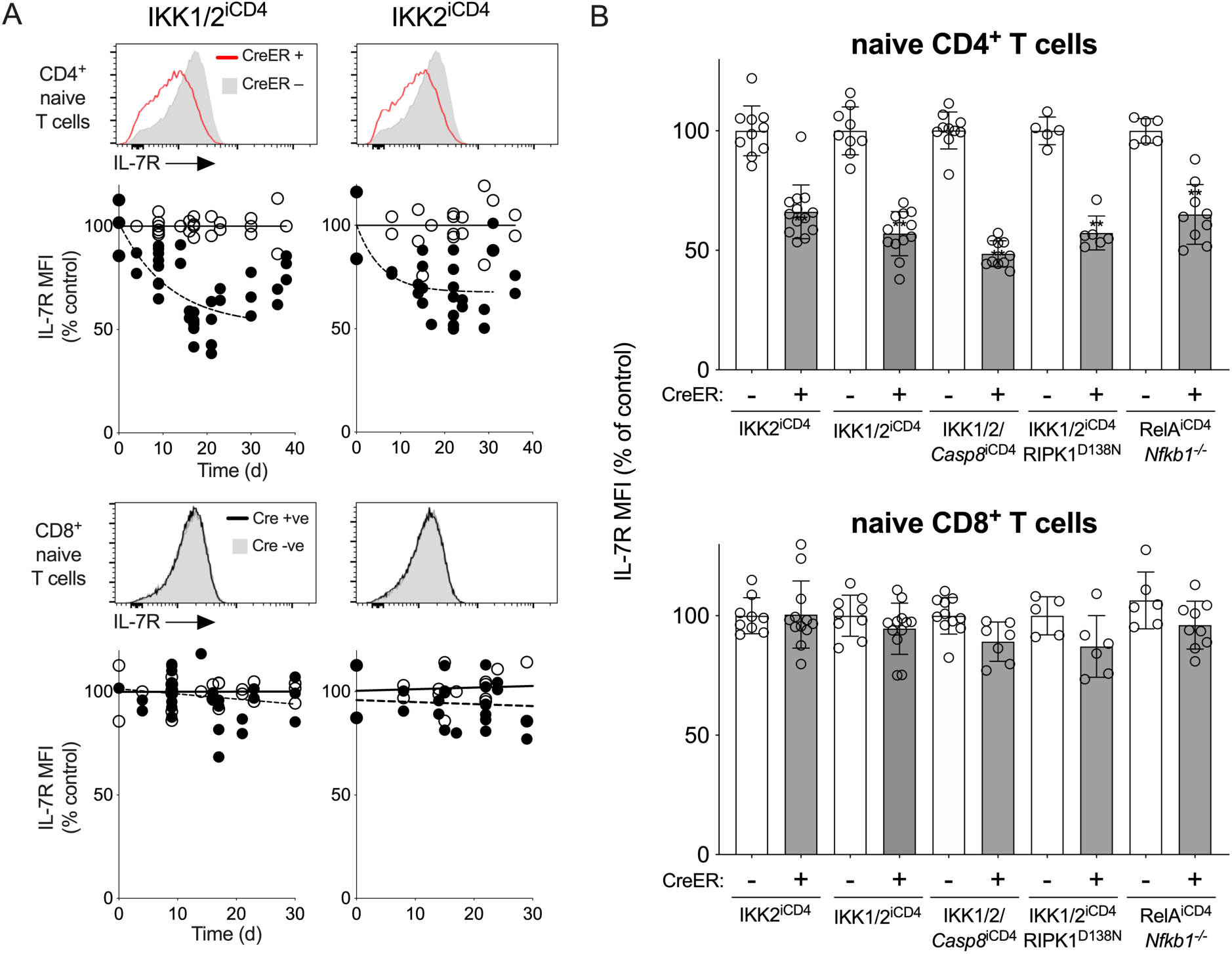
Maintenance of IL-7R expression is dependent upon tonic NF-κB signalling. T cells from groups of IKK2^iCD4^, IKK1/2^iCD4^, IKK1/2/Casp8^iCD4^, IKK1/2^iCD4^ RIPK1^D138N^, RelA^iCD4^*Nfkb1^-/-^* mice, together with *Cd4^CreER^* -ve littermate controls, described in figures 2, 5 and 6 were analysed for expression of IL-7R ∼3 weeks after start of tamoxifen treatment. (A) Histograms are of IL-7R expression by CD4^+^ CD44^lo^CD25^-^ and CD8^+^ CD44^lo^CD25^-^ naive T cells from CreERT+ (lines) or CeERT-(grey fills) IKK2^iCD4^ and IKK1/2^iCD4^ mice from d17 following start of tamoxifen treatment. Scatter plots show normalised IL-7R MFI by CreERT+ (filled circles) or CeERT-(empty circles) IKK2^iCD4^ and IKK1/2^iCD4^ mice at various times following onset of tamoxifen treatment. IL-7R MFI was normalised against the mean MFI of CeERT-littermates analysed at the same time. (B) Bar charts shows normalised IL-7R MFI, of naive CD4^+^ and CD8^+^ T cells from the indicated strains, analysed between 17 and 22 days after onset of tamoxifen treatment. Circles indicate normalised IL-7R MFI as a % of controls, of T cells from individual mice.

### Discussion

Although IKK was first characterised as the critical regulator of NF-κB activation through its ability to target inhibitory IκB proteins for degradation by their phosphorylation, a role in directly repressing cell death triggers has more recently been recognised, mediated by direct phosphorylation, and thereby repression, of RIPK1 (Dondelinger et al., 2019; Dondelinger et al., 2015). Survival of developing thymocytes depends exclusively on this repressive activity upon extrinsic cell death pathways and not on NF-κB activation(Webb et al., 2019). In this study, we addressed for the first time the role of IKK signalling in the maintenance of mature peripheral naive T cells. We show that naive T cells are subject to tonic IKK signalling, and like their thymic precursors, a loss of IKK expression resulted in a profound and rapid collapse in peripheral T cell numbers. However, our new results reveal contrasting roles for NF-κB and extrinsic cell death signalling for survival of peripheral naive T cells compared with their thymic progenitors. Even though naive T cells exhibit heightened sensitivity to RIPK1 dependent cell death in response to TNF, compared with thymocytes, this mechanism only partially accounted for the loss of T cells following IKK1/2 ablation. Instead, activation of NF-κB by IKK also appears to be an important mechanism promoting longterm survival of peripheral naive T cells. The extent of RIPK1 independent cell death observed in RIPK1^D138N^ IKK1/2^iCD4^ mice closely matched that following RELA ablation in RelA^iCD4^ mice. The most parsimonious conclusion is that RIPK1 independent and RELA dependent cell death processes in these two scenarios are one and the same.

IL-7 is well recognised as one of the most crucial survival factors for peripheral T cells and regulation of IL7R expression is a critical point of control of this pathway (Barata et al., 2019). Our results show that tonic signalling from TNFRSF receptors is also critical for naive T cell survival. Induction of IL7R in thymocytes depends on NF-κB(Silva et al., 2014). Our data shows that maintenance of IL7R expression by mature naive T cells continues to rely upon continued NF-κB expression and tonic IKK signalling, since loss of RELA or upstream IKK resulted in a similar loss of IL7R. Therefore, maintenance of IL-7R abundance is likely to be one important mechanism by which NF-κB signalling promotes naive T cell survival, thereby placing TNFRSF signalling upstream of IL-7 signalling in controlling long term survival of naive T cells. The loss of IL7R following peripheral IKK1/2 ablation did not, however, appear as substantial as the defect in IL-7R induction observed in constitutive RELA/NFKB1/cREL deficient mice(Webb et al., 2019). This may reflect the complex nature of the experiment, in which tamoxifen induced gene deletion over five days may create asynchronous waves of gene deleted T cells that are lost to varying extents by the time of readout. There may also be counter selection against cells with the lowest IL7R levels in the three weeks following gene ablation. Loss of NF-κB signalling in T cells results in global changes in gene transcription(Webb et al., 2019), and it is possible that the reduction in IL7R is not solely responsible for the failure of naive CD4^+^ T cells to survive following inducible RELA deletion. Identifying other functionally important NF-κB targets for naive T cell survival will be an important area future research.

The stark loss of mature SP thymocytes to TNF induced cell death observed when IKK1/2 is deleted in the T cell lineage (Webb et al., 2016) reveals fundamental developmental changes in extrinsic cell death signalling that occur following positive selection of T cells in the thymus. Induction of RIPK1 expression as SP thymocytes mature, renders them highly susceptible to extrinsic cell death that necessitates the repressive activity of the IKK complex. Our new data show that engagement and maturation of extrinsic cell death pathways continues further as new naive T cells complete maturation. Careful comparison of TNF reactivity by different T cell populations revealed that peripheral naive T cells exhibit higher levels of RIPK1 activity in response to TNF stimulation than do thymocytes, and that this is also greater in CD8^+^ lineage cells than in CD4^+^ T cells. Indeed, inhibition of IKK2 alone is sufficient to render naive T cells susceptible to some TNF induced cell death in vitro. In vivo, IKK2 deletion resulted in modest cell loss, further suggesting that repression of RIPK1 in mature T cells requires greater IKK activity than is required in thymocytes. Although many cell types exhibit similar requirements for IKK expression for normal tissue homeostasis, T cells are somewhat unique in exhibiting such sophisticated developmental regulation of the signalling pathways, suggesting that these pathways serve specific functions at different developmental stages. In addition to sensitising cells to extrinsic cell death, induction of RIPK1 also couples TNFRSF signalling to NF-κB activation in T cells(Webb et al., 2019). The timing of RIPK1 induction in thymocytes, specifically following positive selection, suggests that permitting TNFRSF signalling during positive selection could be problematic, potentially uncoupling DP thymocyte survival from selfMHC restriction of their TCRs. Indeed, another critical T cell survival cue, IL7R, is also lost during positive selection (Barata et al., 2019). In addition, it is also unclear whether the tuning of extrinsic cell death pathways continues further in mature peripheral T cells. T cell activation represents another critical checkpoint in the T cell life course, and the role that IKK regulation of extrinsic cell death pathways might play, or whether the pathways are tuned further, has not been investigated to date. One clue comes from *Casp8* deficient T cells, that only become susceptible to an alternative mode of RIPK1 dependent cell death, necroptosis, following T cell activation(Ch’en et al., 2011), demonstrating at least one facet of extrinsic death pathways that does change.

Finally, a key outstanding question from our studies is the identity of the receptors that trigger IKK and death signalling in mature T cells. We have previously shown that TNFR1 ablation is sufficient to rescue CD4^+^ SP thymocytes and largely, though not completely, rescue CD8^+^ SP thymocyte survival (Webb et al., 2016). Here, we confirmed these results using *Tnf* ablation instead, which phenocopies TNFR1 deletion and excludes an unlikely role for TNFR2 as a death trigger. However, we found that *Casp8* deletion achieved a greater rescue of cell survival from IKK1/2 deletion than TNF or TNFR1. *Casp8* deletion rescued both CD8^+^ SP thymocyte survival to normality in IKK1/2/*Casp8*^ΔCD2^ mice and peripheral CD4^+^ naive T cells following inducible *Cd4^CreERT2^* mediated IKK1/2 deletion. Together, these results suggest the activity of other death inducing TNFRSFs receptors, that first impact cell survival in mature CD8 SP thymocytes, and have a redundant activity along side TNF in peripheral T cells in the absence of IKK. Redundancy between TNFRSF members such as TNFR1, TNFR2, OX40, GITR and CD27 to activate NF-κB in T cells has already been reported (Mahmud et al., 2014; Silva et al., 2014). That cell death of peripheral naive T cells occurs in the absence of IKK and is RIPK1 dependent, suggests the activity of TNFRSF members with death domains that can recruit TRADD, such as DR3 and TRAILR, since E3 ligase mediated assembly of ubiquitin scaffold that occurs downstream of TRADD, is required for recruitment of IKK and RIPK1 (Blanchett et al., 2021). Identifying the full range of triggers for these critical signals to naive T cells is an important future challenge, and underlines the complex and shifting landscape with which NF-κB and extrinsic death pathways regulate development and maintenance of T cells.

### Materials and methods

#### Mice

Mice with conditional alleles of *Ikbkb* (Li et al., 2003)*, Chuk* (Gareus et al., 2007), *Rela* (Steinbrecher et al., 2008), *Casp8* were intercrossed with mice either expressing *Cre* under the control of the human CD2 (*huCD2^iCre^*)(de Boer et al., 2003), mice expressing Cre^ERT2^ under control of *Cd4* expression elements (*Cd4^CreERT^*)(Sledzinska et al., 2013) or *Rosa26* (*Rosa26^CreERT^*) (de Luca et al., 2005), or *Tnf*^-/-^, *Nfkb1*^-/-^, *Rel*^-/-^ mice, or mice with a D138N mutation in *Ripk1* (RIPK1^D138N^)(Newton et al., 2014). *Chuk^fx/fx^ Ikbkb^fx/fx^Rosa26^CreERT^, Chuk^fx/fx^ Ikbkb^fx/fx^ Cd4^CreERT^* (IKK1/2^iCD4^), *Ikbkb^fx/fx^ Cd4^CreERT^* (IKK2^iCD4^), *Chuk^fx/fx^ huCD2^iCre^* (IKK1^ΔCD2^), *Chuk^fx/fx^ Ikbkb^fx/fx^ huCD2^iCre^* (IKK1/2^ΔCD2^), *Chuk^fx/fx^ Ikbkb^fx/fx^ Casp8^fx/fx^ huCD2^iCre^* (IKK1/2/Casp8^ΔCD2^), *Chuk^fx/fx^ Ikbkb^fx/ fx^ Casp8^fx^*^/fx^ *Cd4^CreERT^* (IKK1/2/Casp8^iCD4^), *Chuk^fx/fx^ Ikbkb^fx/fx^ Cd4^CreERT^* RIPK1^D138N^ (IKK1/2^iCD4^ RIPK1^D138N^), *Chuk^fx/fx^ Ikbkb^fx/fx^ Cd4^CreERT^ Tnf^-/-^* (IKK1/2^iCD4^ *Tnf^-/-^*), *Rela^fx/fx^ Cd4^CreERT^ Nfkb1^-/-^* (*Rela^iCD4^ Nfkb1^-/-^*), *Rela^fx/fx^ Cd4^CreERT^ Rel^-/-^* (*Rela^iCD4^ Rel^-/-^*), *Rela^fx/fx^ Rel^fx/fx^ Cd4^CreERT^*(*Rela.Rel^iCD4^*), B6.SJL, (B6.SJL x C57Bl6/J)F1 and *Rag1*^-/-^ strains were bred in the Comparative Biology Unit of the Royal Free UCL campus and at Charles River laboratories, Manston, UK. Cd4^CreERT2^ and Rosa26^CreERT^ activity was induced *in vivo* by i.p. injection of 2mg of tamoxifen in corn oil for five consecutive days. Irradiation chimeras were generated following either sublethal irradiation of *Rag1*^-/-^ hosts with 500rads or lethal irradiation of B6.SJL hosts with 1000rads. Shortly after, mice were injected with 10^7^ depleted bone marrow cells from the induceible strains, and in some experiments, control bone marrow from (B6.SJL x C57Bl6/J)F1 donors, and allowed to reconstitute for eight weeks. Animal experiments were performed according to institutional guidelines and Home Office regulations.

#### Flow cytometry and electronic gating strategies

Flow cytometric analysis was performed with 2-5 x 10^6^ thymocytes, 1-5 x 10^6^ lymph node or spleen cells. Cell concentrations of thymocytes, lymph nodes (superficial cervical, mandibular, axillary, superficial inguinal, and mesenteric chain) and spleen cells were determined with a Scharf Instruments Casy Counter. Cells were incubated with saturating concentrations of antibodies in 100 μl of Dulbecco’s phosphate-buffered saline (PBS) containing bovine serum albumin (BSA, 0.1%) for 1hour at 4°C followed by two washes in PBS-BSA. Panels used the following mAb: EF450-conjugated antibody against CD25(ThermoFisher Scientific), PE-conjugated antibody against CD127 (ThermoFisher Scientific), BV785-conjugated CD44 antibody (Biolegend), BV650-conjugated antibody against CD4 (Biolegend), BUV395-conjugated antibody against CD8 (BD Biosciences), BUV737-conjugated antibody against CD24 (BD Biosciences), PerCP-cy5.5-conjugated antibody against TCR (Tonbo Biosciences). Cell viability was determined using LIVE/ DEAD cell stain kit (Invitrogen Molecular Probes), following the manufacturer’s protocol. multi-color flow cytometric staining was analyzed on a LSRFortessa (Becton Dickinson) instrument, and data analysis and color compensations were performed with FlowJo V10 software (TreeStar). Naive peripheral T cells were identified by gating CD4^+^ or CD8^+^ subsets with TCR^hi^ CD44^lo^ CD25^lo^. Mature CD4^+^ and CD8^+^ SP thymocytes were identified as TCR^hi^CD4^+^CD8^-^HSA^lo^ and TCR^hi^CD4^-^ CD8^+^HSA^lo^ respectively.

#### In vitro culture

Thymocytes and LN T cells were cultured at 37^°^C with 5% CO2 in RPMI-1640 (Gibco, Invitrogen Corporation, CA) supplemented with 10% (v/v) fetal bovine serum (FBS) (Gibco Invitrogen), 0.1% (v/v) 2-mercaptoethanol βME (Sigma Aldrich) and 1% (v/v) penicillin-streptomycin (Gibco Invitrogen) (RPMI-10). Recombinant TNF was supplemented to cultures at 20ng/ml, unless otherwise stated, and was obtained from Peprotech, with PBS used as vehicle. Inhibitors were used at the following concentrations, unless otherwise stated: IKK2 inhibitor BI605906 (IKK2i) (10µM in 0.1% DMSO vehicle), IKK16 (2µM in 0.1% DMSO), Nec1 (10µM in 0.1% DMSO).

#### Immunoblotting

10^6^ purified CD4^+^ T cells were used per condition. Cell extracts were analyzed by NuPage 3-8% Bis-Tris gel (Invitrogen Novex), transferred onto PVDF membrane (Millipore) and immunoblotted with anti-IKK1 (Cell Signalling Technology), anti-IKK2 (Cell Signalling Technology) and tubulin (Cell Signalling Technology). Immunodetection was performed by incubation with horseradish peroxidise-conjugated anti-rabbit (1:5000) (DAKO) and developed by enhanced chemiluminescence (Millipore).

#### Statistics

Statistical analysis, line fitting, regression analysis, and figure preparation were performed using Graphpad Prism 8. Column data compared by unpaired Mann-Witney student’s t test. * p<0.05, ** p<0.01, *** p<0.001, **** p < 0.0001.

## Acknowledgements

We thank UCL Comparative Biology Unit staff for assistance with mouse breeding and maintenance. We thank the following for generously sharing of their mouse strains: Prof Manolis Pasparakis for *Chuk* conditional strain, Prof Michael Karin for *Ikbkb* conditional strain, Prof Vishva Dixit for the RIPK1^D138N^ strain, Prof Albert Baldwin for *Rela* conditional strain, Prof Steve Gerondakis for *Nfkb1*^-/-^ and *cRel*^-/-^ mice and Prof Burkhard Becher for the *Cd4^CreERT^* strain. The authors declare no competing financial interests.

## Funding

The work in the Seddon lab is supported by the Medical Research Council UK under programme codes MR/P011225/1 and MR/N013867/1.

## Author contributions

Conceptualization: FC, BS

Methodology: FC, SL, AB, LW, BS, FI

Investigation: FC, SL, AB, LW, FI

Visualization: FC, BS

Funding acquisition: BS

Project administration: FC, BS

Supervision: BS

Writing – original draft: BS

Writing – review & editing: BS, SL

---

**Figure S1.**
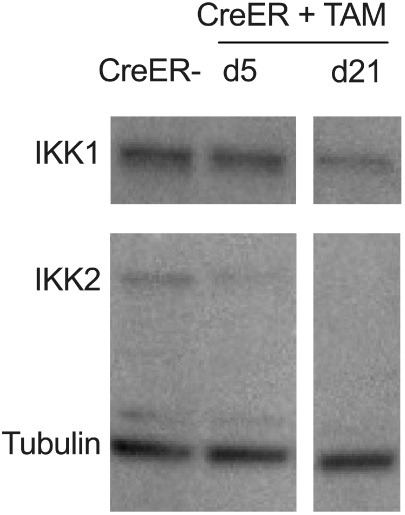
IKK1 and IKK2 protein expression following tamxoifen induced gene deletion. IKK1/2^iCD4^ and Cre-litter mates (WT) were treated with tamoxifen for five days. At d5 and d21, total CD4^+^ T cells were isolated by magnetic separation to >96% purity. Cytosolic cell extracts from 10^6^ purified T cells were analysed by immunoblotting for expression of IKK1, IKK2 and tubulin. Data are representative of two independent experiments.

**Figure S2.**
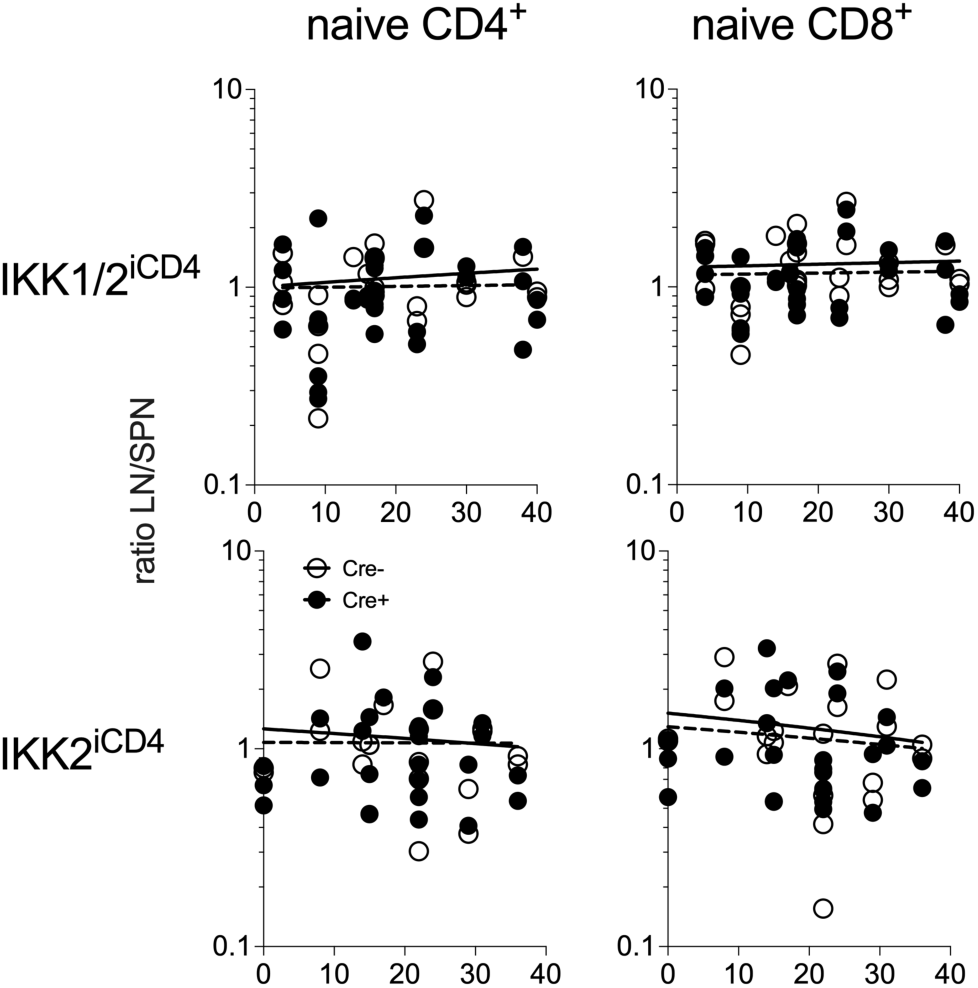
Distribution of naive T cells between lymph node and spleen of tamoxifen treated mice. Scatter plots show the distribution of naive CD4^+^ and CD8^+^ T cells between lymph nodes (superficial cervical, mandibular, axillary, superficial inguinal, and mesenteric chain) and spleen, recovered from experimental tamoxifen treated IKK1/2^iCD4^ and IKK2^iCD4^ mice described in figure 2. Scatter plot shows ratio of naive T cells in lymph node : spleen of individual mice. Line plots are of best fit linear regression, with no significant deviation in slope from 0, and approximately equal distribution (i.e. ratio of 1) between lymph node and spleen of both strains.

**Figure S3.**
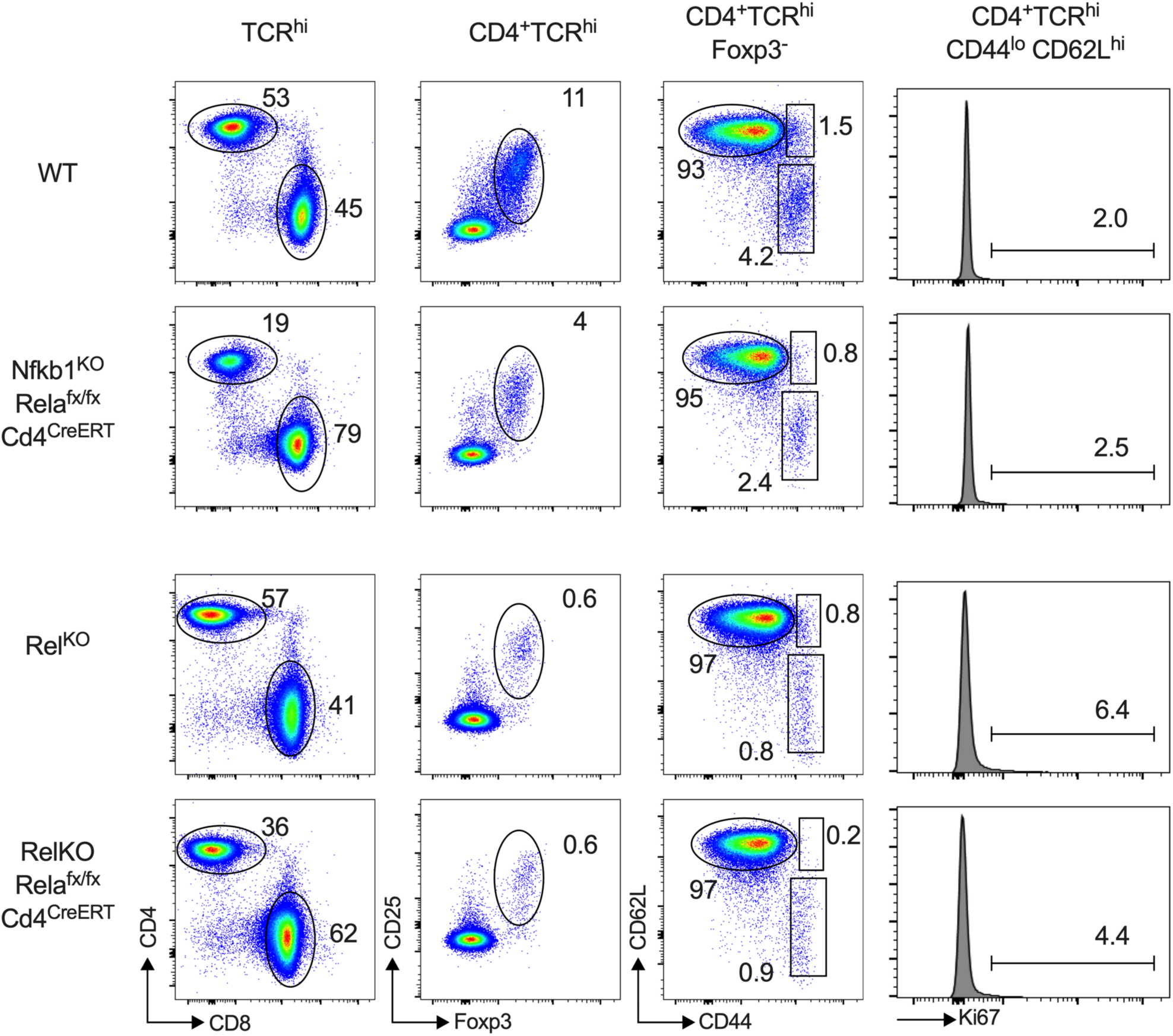
CD4+ Naive T cells loss following RelA deletion is independent of Treg. Flow analysis of tamoxifen treated mice from the indicated strains from experiments described in main figure 6.

## Notes

### Competing Interest Statement

The authors have declared no competing interest.

### Summary of Updates

Revised version for further submission

## Bibliography

Annibaldi, A., and P. Meier. 2018. Checkpoints in TNF-Induced Cell Death: Implications in Inflammation and Cancer. Trends Mol Med 24:49–65.

Barata, J.T., S.K. Durum, and B. Seddon. 2019. Flip the coin: IL-7 and IL-7R in health and disease. Nat Immunol 20:1584–1593.

Blanchett, S., I. Boal-Carvalho, S. Layzell, and B. Seddon. 2021. NF-kappaB and Extrinsic Cell Death Pathways - Entwined Do-or-Die Decisions for T cells. Trends Immunol 42:76–88.

Bonizzi, G., and M. Karin. 2004. The two NF-kappaB activation pathways and their role in innate and adaptive immunity. Trends Immunol 25:280–288.

Ch’en, I.L., J.S. Tsau, J.D. Molkentin, M. Komatsu, and S.M. Hedrick. 2011. Mechanisms of necroptosis in T cells. Journal of Experimental Medicine 208:633–641.

Clark, K., M. Peggie, L. Plater, R.J. Sorcek, E.R. Young, J.B. Madwed, J. Hough, E.G. McIver, and P. Cohen. 2011. Novel cross-talk within the IKK family controls innate immunity. Biochem J 434:93–104.

de Boer, J., A. Williams, G. Skavdis, N. Harker, M. Coles, M. Tolaini, T. Norton, K. Williams, K. Roderick, A.J. Potocnik, and D. Kioussis. 2003. Transgenic mice with hematopoietic and lymphoid specific expression of Cre. Eur J Immunol 33:314–325.

de Luca, C., T.J. Kowalski, Y. Zhang, J.K. Elmquist, C. Lee, M.W. Kilimann, T. Ludwig, S.M. Liu, and S.C. Chua, Jr. 2005. Complete rescue of obesity, diabetes, and infertility in db/db mice by neuron-specific LEPR-B transgenes. J Clin Invest 115:3484–3493.

Dondelinger, Y., M.A. Aguileta, V. Goossens, C. Dubuisson, S. Grootjans, E. Dejardin, P. Vandenabeele, and M.J. Bertrand. 2013. RIPK3 contributes to TNFR1-mediated RIPK1 kinase-dependent apoptosis in conditions of cIAP1/2 depletion or TAK1 kinase inhibition. Cell Death Differ 20:1381–1392.

Dondelinger, Y., T. Delanghe, D. Priem, M.A. Wynosky-Dolfi, D. Sorobetea, D. Rojas-Rivera, P. Giansanti, R. Roelandt, J. Gropengiesser, K. Ruckdeschel, S.N. Savvides, A.J.R. Heck, P. Vandenabeele, I.E. Brodsky, and M.J.M. Bertrand. 2019. Serine 25 phosphorylation inhibits RIPK1 kinase-dependent cell death in models of infection and inflammation. Nat Commun 10:1729.

Dondelinger, Y., S. Jouan-Lanhouet, T. Divert, E. Theatre, J. Bertin, P.J. Gough, P. Giansanti, A.J. Heck, E. Dejardin, P. Vandenabeele, and M.J. Bertrand. 2015. NF-kappaB-Independent Role of IKKalpha/IKKbeta in Preventing RIPK1 Kinase-Dependent Apoptotic and Necroptotic Cell Death during TNF Signaling. Mol Cell 60:63–76.

Dondelinger, Y., P. Vandenabeele, and M.J. Bertrand. 2016. Regulation of RIPK1’s cell death function by phosphorylation. Cell Cycle 15:5–6.

Egawa, T., B. Albrecht, B. Favier, M.J. Sunshine, K. Mirchandani, W. O’Brien, M. Thome, and D.R. Littman. 2003. Requirement for CARMA1 in antigen receptor-induced NF-kappa B activation and lymphocyte proliferation. Curr Biol 13:1252–1258.

Gareus, R., M. Huth, B. Breiden, A. Nenci, N. Rosch, I. Haase, W. Bloch, K. Sandhoff, and M. Pasparakis. 2007. Normal epidermal differentiation but impaired skin-barrier formation upon keratinocyte-restricted IKK1 ablation. Nat Cell Biol 9:461–469.

Hara, H., T. Wada, C. Bakal, I. Kozieradzki, S. Suzuki, N. Suzuki, M. Nghiem, E.K. Griffiths, C. Krawczyk, B. Bauer, F. D’Acquisto, S. Ghosh, W.C. Yeh, G. Baier, R. Rottapel, and J.M. Penninger. 2003. The MAGUK family protein CARD11 is essential for lymphocyte activation. Immunity 18:763–775.

Li, Q., Q. Lu, J.Y. Hwang, D. Buscher, K.F. Lee, J.C. Izpisua-Belmonte, and I.M. Verma. 1999. IKK1-deficient mice exhibit abnormal development of skin and skeleton. Genes Dev 13:1322–1328.

Li, Z.W., S.A. Omori, T. Labuda, M. Karin, and R.C. Rickert. 2003. IKK beta is required for peripheral B cell survival and proliferation. J Immunol 170:4630–4637.

Liu, H.H., M. Xie, M.D. Schneider, and Z.J. Chen. 2006. Essential role of TAK1 in thymocyte development and activation. Proceedings of the National Academy of Sciences of the United States of America 103:11677–11682.

Mahmud, S.A., L.S. Manlove, H.M. Schmitz, Y. Xing, Y. Wang, D.L. Owen, J.M. Schenkel, J.S. Boomer, J.M. Green, H. Yagita, H. Chi, K.A. Hogquist, and M.A. Farrar. 2014. Costimulation via the tumor-necrosis factor receptor superfamily couples TCR signal strength to the thymic differentiation of regulatory T cells. Nature Immunology 15:473–481.

Miller, M.L., M. Mashayekhi, L. Chen, P. Zhou, X. Liu, M. Michelotti, N. Tramontini Gunn, S. Powers, X. Zhu, C. Evaristo, M.L. Alegre, and L.L. Molinero. 2014. Basal NF-kappaB controls IL-7 responsiveness of quiescent naive T cells. Proc Natl Acad Sci U S A 111:7397–7402.

Newton, K., D.L. Dugger, K.E. Wickliffe, N. Kapoor, M.C. de Almagro, D. Vucic, L. Komuves, R.E. Ferrando, D.M. French, J. Webster, M. Roose-Girma, S. Warming, and V.M. Dixit. 2014. Activity of protein kinase RIPK3 determines whether cells die by necroptosis or apoptosis. Science 343:1357–1360.

Oh, H., Y. Grinberg-Bleyer, W. Liao, D. Maloney, P. Wang, Z. Wu, J. Wang, D.M. Bhatt, N. Heise, R.M. Schmid, M.S. Hayden, U. Klein, R. Rabadan, and S. Ghosh. 2017. An NF-kappaB Transcription-Factor-Dependent Lineage-Specific Transcriptional Program Promotes Regulatory T Cell Identity and Function. Immunity 47:450–465 e455.

Pasparakis, M., G. Courtois, M. Hafner, M. Schmidt-Supprian, A. Nenci, A. Toksoy, M. Krampert, M. Goebeler, R. Gillitzer, A. Israel, T. Krieg, K. Rajewsky, and I. Haase. 2002. TNF-mediated inflammatory skin disease in mice with epidermis-specific deletion of IKK2. Nature 417:861–866.

Ruefli-Brasse, A.A., D.M. French, and V.M. Dixit. 2003. Regulation of NF-kappaB-dependent lymphocyte activation and development by paracaspase. Science 302:1581–1584.

Ruland, J., G.S. Duncan, A. Elia, I. del Barco Barrantes, L. Nguyen, S. Plyte, D.G. Millar, D. Bouchard, A. Wakeham, P.S. Ohashi, and T.W. Mak. 2001. Bcl10 is a positive regulator of antigen receptor-induced activation of NF-kappaB and neural tube closure. Cell 104:33–42.

Ruland, J., G.S. Duncan, A. Wakeham, and T.W. Mak. 2003. Differential requirement for Malt1 in T and B cell antigen receptor signaling. Immunity 19:749–758.

Schmidt-Supprian, M., W. Bloch, G. Courtois, K. Addicks, A. Israel, K. Rajewsky, and M. Pasparakis. 2000. NEMO/IKK gamma-deficient mice model incontinentia pigmenti. Mol Cell 5:981–992.

Schmidt-Supprian, M., G. Courtois, J. Tian, A.J. Coyle, A. Israel, K. Rajewsky, and M. Pasparakis. 2003. Mature T cells depend on signaling through the IKK complex. Immunity 19:377–389.

Schmidt-Supprian, M., J. Tian, E.P. Grant, M. Pasparakis, R. Maehr, H. Ovaa, H.L. Ploegh, A.J. Coyle, and K. Rajewsky. 2004. Differential dependence of CD4+CD25+ regulatory and natural killer-like T cells on signals leading to NF-kappaB activation. Proc Natl Acad Sci U S A 101:4566–4571.

Silva, A., G. Cornish, S.C. Ley, and B. Seddon. 2014. NF-kappaB signaling mediates homeostatic maturation of new T cells. Proc Natl Acad Sci U S A 111:E846–855.

Sledzinska, A., S. Hemmers, F. Mair, O. Gorka, J. Ruland, L. Fairbairn, A. Nissler, W. Muller, A. Waisman, B. Becher, and T. Buch. 2013. TGF-beta signalling is required for CD4(+) T cell homeostasis but dispensable for regulatory T cell function. PLoS Biol 11:e1001674.

Srinivas, S., T. Watanabe, C.S. Lin, C.M. William, Y. Tanabe, T.M. Jessell, and F. Costantini. 2001. Cre reporter strains produced by targeted insertion of EYFP and ECFP into the ROSA26 locus. BMC Dev Biol 1:4.

Steinbrecher, K.A., E. Harmel-Laws, R. Sitcheran, and A.S. Baldwin. 2008. Loss of epithelial RelA results in deregulated intestinal proliferative/apoptotic homeostasis and susceptibility to inflammation. J Immunol 180:2588–2599.

Tanaka, M., M.E. Fuentes, K. Yamaguchi, M.H. Durnin, S.A. Dalrymple, K.L. Hardy, and D.V. Goeddel. 1999. Embryonic lethality, liver degeneration, and impaired NF-kappa B activation in IKK-beta-deficient mice. Immunity 10:421–429.

Ting, A.T., and M.J.M. Bertrand. 2016. More to Life than NF-kappaB in TNFR1 Signaling. Trends Immunol 37:535–545.

Tummers, B., and D.R. Green. 2017. Caspase-8: regulating life and death. Immunol Rev 277:76–89.

Vandenabeele, P., W. Declercq, F. Van Herreweghe, and T. Vanden Berghe. 2010. The role of the kinases RIP1 and RIP3 in TNF-induced necrosis. Sci Signal 3:re4.

Wan, Y.Y., H. Chi, M. Xie, M.D. Schneider, and R.A. Flavell. 2006. The kinase TAK1 integrates antigen and cytokine receptor signaling for T cell development, survival and function. Nature immunology 7:851–858.

Wang, L., F. Du, and X. Wang. 2008. TNF-alpha induces two distinct caspase-8 activation pathways. Cell 133:693–703.

Webb, L.V., A. Barbarulo, J. Huysentruyt, T. Vanden Berghe, N. Takahashi, S. Ley, P. Vandenabeele, and B. Seddon. 2019. Survival of Single Positive Thymocytes Depends upon Developmental Control of RIPK1 Kinase Signaling by the IKK Complex Independent of NF-kappaB. Immunity 50:348–361 e344.

Webb, L.V., S.C. Ley, and B. Seddon. 2016. TNF activation of NF-kappaB is essential for development of single-positive thymocytes. J Exp Med 213:1399–1407.

Xing, Y., X. Wang, S.C. Jameson, and K.A. Hogquist. 2016. Late stages of T cell maturation in the thymus involve NF-κB and tonic type I interferon signaling. Nature Immunology 17:565–573.

Zaph, C., A.E. Troy, B.C. Taylor, L.D. Berman-Booty, K.J. Guild, Y. Du, E.A. Yost, A.D. Gruber, M.J. May, F.R. Greten, L. Eckmann, M. Karin, and D. Artis. 2007. Epithelial-cell-intrinsic IKK-beta expression regulates intestinal immune homeostasis. Nature 446:552–556.

Zheng, Y., M. Vig, J. Lyons, L. Van Parijs, and A.A. Beg. 2003. Combined deficiency of p50 and cRel in CD4+ T cells reveals an essential requirement for nuclear factor kappaB in regulating mature T cell survival and in vivo function. The Journal of experimental medicine 197:861–874.

