## Supplementary material for "IKK controls naive T cell survival by repressing RIPK1 dependent apoptosis and activation of NF-κB": Supp Figs S1-S3

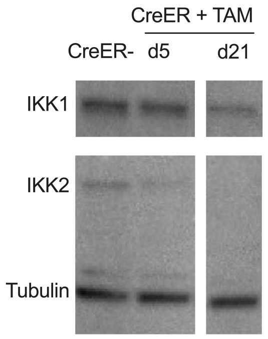


**Figure S1 - IKK1 and IKK2 protein expression following tamxoifen induced gene deletion.** IKK1/2$^{iCD4}$ and Cre- litter mates (WT) were treated with tamoxifen for five days. At d5 and d21, total CD4$^{+}$ T cells were isolated by magnetic separation to >96% purity. Cytosolic cell extracts from $10^6$ purified T cells were analysed by immunoblotting for expression of IKK1, IKK2 and tubulin. Data are representative of two independent experiments.

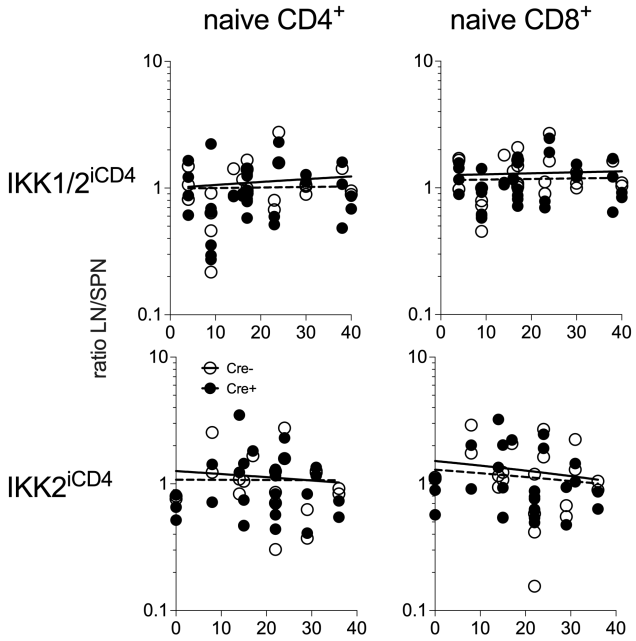


**Figure S2 - Distribution of naive T cells between lymph node and spleen of tamoxifen treated mice.** Scatter plots show the distribution of naive CD4+ and CD8+ T cells between lymph nodes (superficial cervical, mandibular, axillary, superficial inguinal, and mesenteric chain) and spleen, recovered from experimental tamoxifen treated IKK1/2$^{iCD4}$ and IKK2$^{iCD4}$ mice described in figure 2. Scatter plot shows ratio of naive T cells in lymph node : spleen of individual mice. Line plots are of best fit linear regression, with no significant deviation in slope from 0, and approximately equal distribution (i.e. ratio of 1) between lymph node and spleen of both strains.

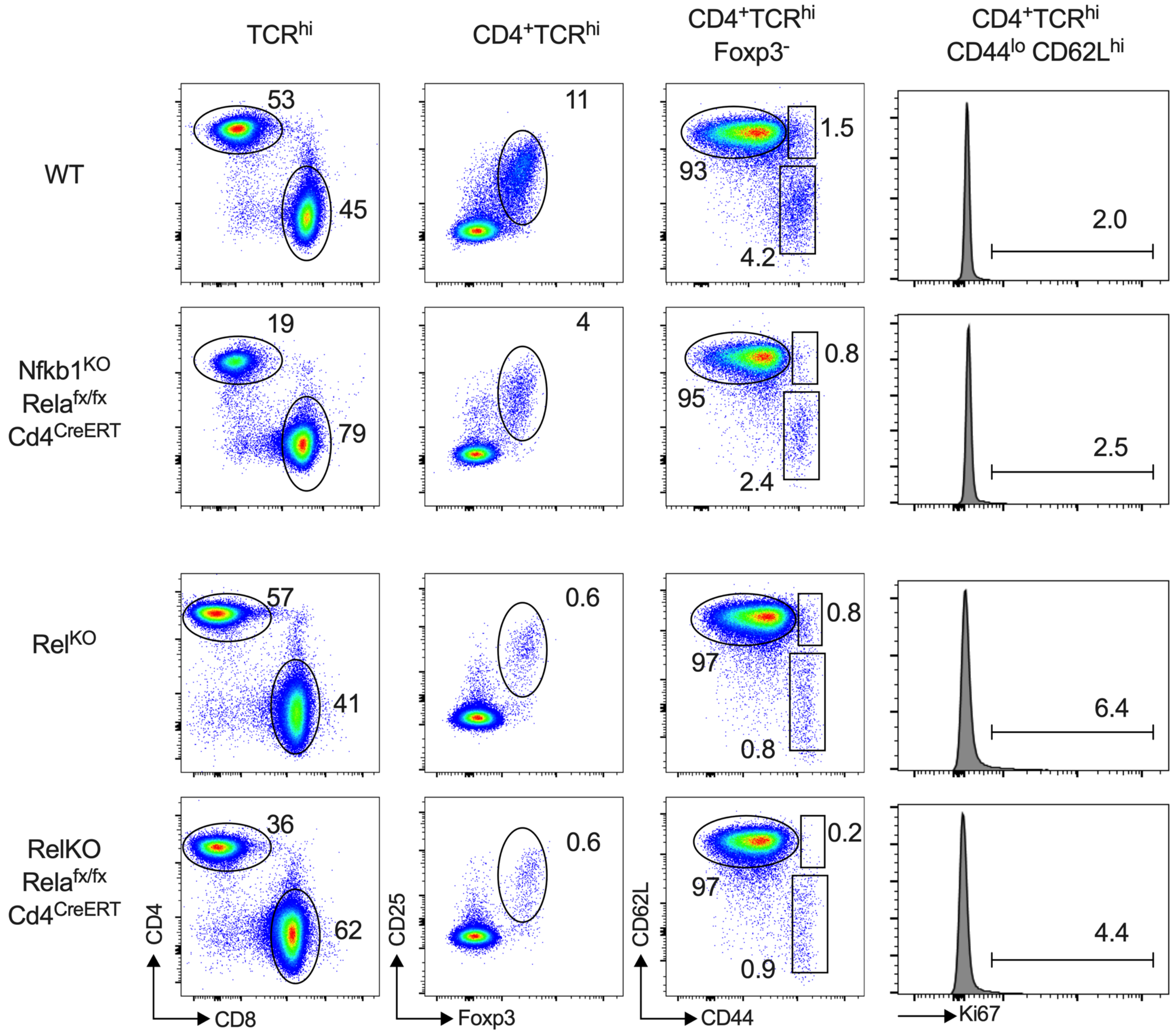


**Figure S3 - CD4+ Naive T cells loss following RelA deletion is independent of Treg.** Flow analysis of tamoxifen treated mice from the indicated strains from experiments described in main figure 6.
